# AB12PHYLO: an integrated pipeline for Maximum Likelihood phylogenetic inference from ABI trace data

**DOI:** 10.1101/2021.03.01.433007

**Authors:** Leo Kaindl, Corinn Small, Remco Stam

**Affiliations:** Chair of Phytopathology, School of Life Sciences Weihenstephan, Technical University of Munich, Freising, Germany

**Keywords:** Barcode sequencing ITS, EF1, Phylogenetic reconstruction, Sanger sequencing, Automation

## Abstract

Multi-gene phylogenies constructed from multiplexed and Sanger sequencing data are regularly used in mycology and other disciplines as a cost-effective way of species identification and as a first means to investigate genetic diversity samples.

Today, a number of tools exist for each of the steps in this analysis, including quality control and trimming, the generation of a multiple sequence alignment (MSA), extraction of informative sites, and the construction of the final phylogenetic tree. A BLAST search in a reference database is often performed to identify sequences of type specimens to compare the samples with in the phylogeny. Made over the past decades, these tools are all independent from and often not perfectly adapted to one another.

We present AB12PHYLO, an integrated pipeline that can perform all necessary steps from reading in raw Sanger sequencing data through visualizing and editing phylogenies. In addition, AB12PHYLO can calculate basic summary statistics for each gene in the phylogeny.

AB12PHYLO is designed as a wrapper of several open access and commonly used tools for each of the intermediate stages, and intended to simplify the phylogenetic pipeline while still allowing a high degree of access. It comes as a command-line version for the highest reproducibility and an intuitive graphical user interface (GUI) for easy adoption by IT-agnostic end-users. The use of AB12PHYLO significantly reduces the hands-on working time for these analyses.

## Main text

Multi-gene phylogenies obtained through Sanger sequencing are a cost-effective and fast way method to aid species identification of fungal samples collected in the field, get the first insight into their genetic diversity, and as a means to select subsets of samples for more expensive and labor-intensive whole genome sequencing approaches. Such analyses often use barcode sequences, specific genic or intergenic fragments of well-defined genes, that have been widely used over the past decades. Examples are regions of the Internally Transcribed Spacer (ITS) (White et al. 1990), Elongation Factor 1 alpha (EF1) (Carbone and Kohn 1999) or RNA polymerase II subunit (RBP2) (Liu et al. 1999). These genes have been sequenced for a large number of type specimens, and sequence comparison of the samples in question with stored type specimens either through direct local alignments or database searches such as NCBI-BLAST (Johnson et al. 2008), can help confirm species identity. Often, sequence data of a single barcode gene is not sufficient to specifically determine fungal identity on species level, whereas a combination of three or more barcodes can reliably determine which species the sample belongs to (see e.g. Woudenberg et al. 2015). In another example, construction of multi-gene phylogenies formed the basis for phylogenetic reclassification: the fungal plant pathogen genus *Ulocladium* appears morphologically different from the genus *Alternaria*, but multi-gene phylogenies did not result in monophyletic clades, suggesting to rename the *Ulocladium* spp, which now fall under the broader *Alternaria* genus (Woudenberg et al. 2013). The method is also in use to get better insights into pathogens in the field: Two recent studies used multi-gene phylogenetic analyses to confirm the nature and relationship of the pathogens *A. alternata* and *A. solani* in potato fields in Wisconsin or similarly in Brazil (Adhikari et al. 2020; Ding et al. 2020). Other recent studies used the method to identify and compare *Colletotrichum* spp. on tea (Orrock et al. 2019), strawberry (Chen et al. 2019), and a variety of hosts (He et al. 2019), to re-assess the taxonomic classification of *Mycosphaerella* spp on persimmon.(Hassan and Chang 2018) or to get first insights in the diversity of Phytophthora spp. in the amazon forest (Legeay et al. 2020).

The analyses presented above often involve manual inspection of the sequence quality, followed by manual data trimming. Some tools exist that automate sequence file inspection to a certain extent (see e.g. Singh and Bhatia 2016; Rausch et al. 2020), yet these tools do not help the user with the subsequent steps, such as alignment with reference sequences or phylogenetic reconstruction, Whereas such steps often require additional hands-on work as well, if only to prepare the output of one tool as input for the next. Manual editing of input and output files would also be the case when using popular web-based phylogeny tools like NGPhylogeny.fr (Lemoine et al. 2019). All manual processing slows down analysis and hampers reproducibility in general, as many parameters or small conversion steps are often not properly recorded. In order to speed up data analyses of this kind and increase their reproducibility, we constructed a fully customizable pipeline which we call AB12PHYLO.

AB12PHYLO is developed as a Python 3 package around widely-used open source tools. It takes raw ab1 (ABI) files. Additionally, the user can provide a template specifying the corresponding sample names, which can be formatted in a 96-well plate format, to represent the way the samples are often loaded for sequencing. When no sample template is specified, AB12PHYLO uses regular expressions that the user can modify to search for and extract the file and gene names.

Its command-line version is assembled from the following three parts (with eight main steps: Part A - Sequence assessment: i) File input: After the command line is supplemented with default configurations, the tables mapping plate coordinates to sample IDs are read to memory. Ab1 trace files are read using Biopython Bio.SeqIO (Cock et al. 2009), matched to their original sample ID and gene, and passed to quality control. Reference sequences are saved to the respective per-gene dataset. ii) Quality control was modeled after SeqTrace (Stucky 2012): Read ends are trimmed until a user-defined proportion of characters in the chromatogram have a phred quality score at or above another user-defined threshold, with 8 / 10 and 30 the pre-set default values. End trimming can discard reads. Consecutive stretches of characters with a score below the phred threshold will be replaced by an equal-length stretch of unknown N characters if they are longer than the last user-definable limit in trace processing; pre-set at 5. Reverse reads are replaced by their reverse complement. Part B – Sequence alignment: iii) Edited sequences are passed to a multiple sequence alignment tool in per-gene datasets. AB12PHYLO is able to interface with local installations of MAFFT (Katoh et al. 2002), Clustal Omega (Sievers et al. 2011) or MUSCLE (Edgar 2004); or an EMBL-EBI online service for any of them (Notredame et al. 2000) at https://www.ebi.ac.uk/Tools/msa.iv) The alignments are trimmed with Gblocks (Castresana 2000). Requirements for a conserved site can be set at four different levels, from 90% identiy to the most relaxed permissible parameters, and a fifth option skipping trimming entirely. The per-gene MSAs are concatenated into a supermatrix alignment. v) A BLAST similarity search of data from the first gene in the analysis is carried out to identify source species. If this search is to be run locally, AB12PHYLO employs BLAST+ (Camacho et al. 2009), which will download, update or check a user-defined database before searching it. Per default, AB12PHYLO will query the NCBI nucleotide database for sequences not found in the local database with Biopython Bio.Blast (Cock et al. 2009), and BLAST can also be run entirely via the public NCBI BLAST API, but this approach is not suitable for large datasets. Two more directly related options are available: Skip BLAST altogether, or parse one or several XML files from a previous analysis or a web BLAST. Part C – Phylogenies: vi) By default, a maximum likelihood (ML) tree is inferred from the concatenated alignment with RAxML-NG (Kozlov et al. 2019). While the evolutionary model is pre-set to GTR+ Γ and the numbers of ML tree searches to 10 with random or parsimony starting trees each, these parameters can be user-defined. Also, the number of parallel threads can be limited. Alternatively, trees can be inferred using IQ-tree (Minh et al. 2020), which allows automated model selection. Moreover, IQ-tree can also be executed in the windows version or AB12PHYLO. vii) With RaxML-NG or IQ-tree, bootstrap replicates are generated from the best ML phylogeny found in the previous step. FBP and TBE support values for the best ML tree are computed from the bootstrap trees constructed in parallel threads. viii) Output: The generated phylogeny is plotted with Toytree (Eaton 2020) and shown alongside other results in an HTML results page. A CGI script allows interactive searching of taxa and selecting populations while computing diversity statistics and the Tajima’s D neutrality test.An overview of the main features of AB12PHYLO is shown in Figure 1. A more detailed model of the command-line AB12PHYLO program flow is shown in Figure S1.

**Figure 1.**
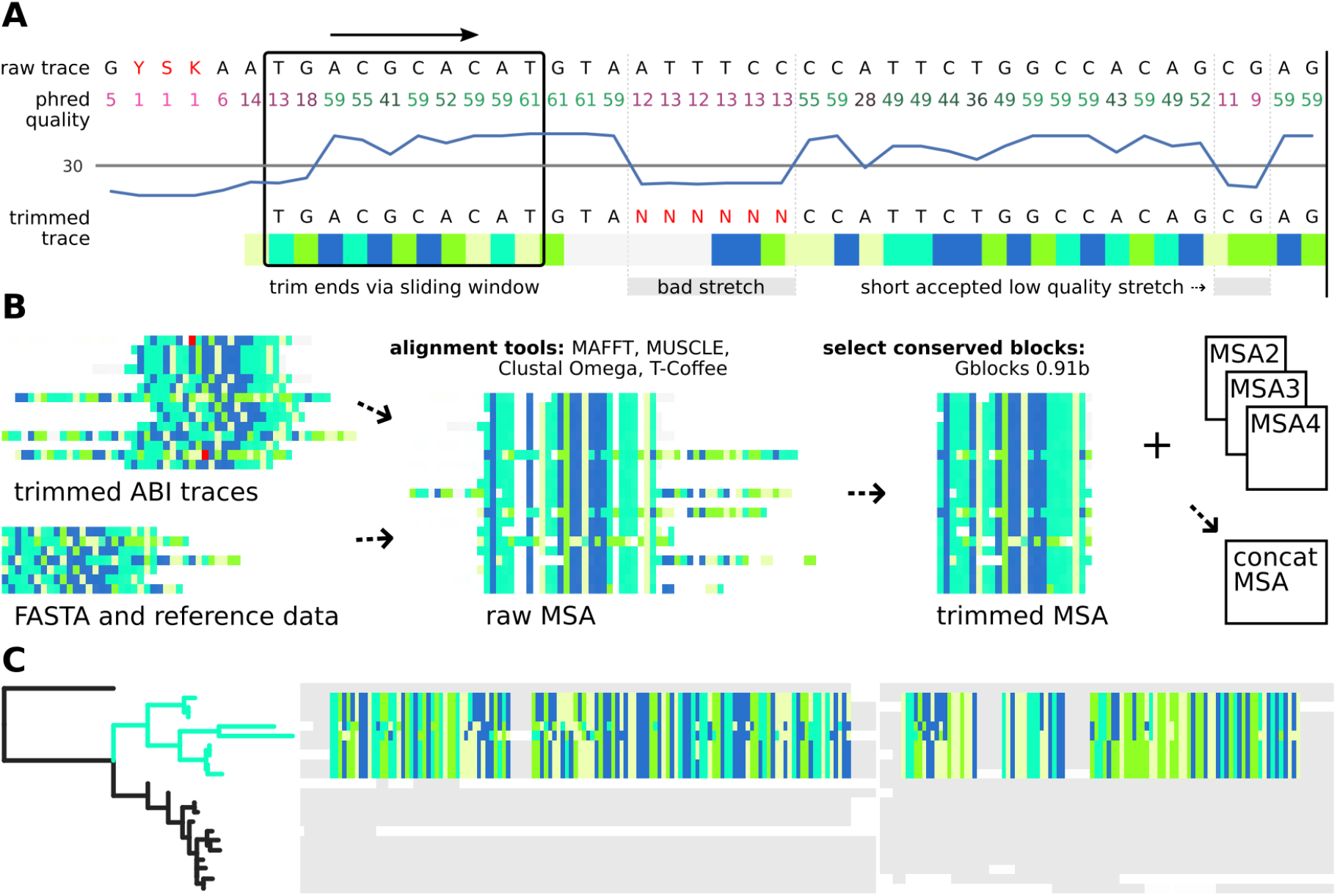
**A:** Sequence data is extracted from ABI trace files using a customisable quality control: Sequence ends are trimmed with a sliding window until a certain number ( *8 out of 10 by default*) of bases reach the minimal accepted phred quality score (*between 0 and 60, 30 by default*). Bases with low phred quality are replaced by N only if they form a consecutive stretch that is longer than a certain threshold (*5 by default*). **B:** Samples missing for a single locus are discarded for all genes. Trimmed traces as well as reference and FASTA sequences are aligned into single-gene Multiple Sequence Alignments (MSAs), which are then each trimmed to a user-defined level conserved positions using Gblocks 0.91b. For multi-gene analyses, the single-gene MSAs are then concatenated into a multi-gene MSA, which is used for ML tree inference. Trees are re-constructed using either RAxML-NG or IQ-Tree 2, with only the latter one available for Windows. **C:** AB12PHYLO allows editing of the resulting tree and selection of taxa by label matching, shared ancestry or manual picking. For these selected sub-populations, basic population genetics neutrality and diversity metrics are calculated from the conserved MSA positions only, with adjustable tolerance of gaps and unknown characters. The graphical ab12phylo is both less cumbersome and more capable for these applications; the wiki pages (ab12phylo, ab12phylo-cmd) have more details.The GUI version of AB12PHYLO implements the same process while giving users direct control over each step: visualizations of sequence trimming and MSAs allow immediate identification of out-of-register samples, and carefully balanced MSA trimming to prevent both signal loss and trimming artifacts. Furthermore, the graphical AB12PHYLO enables comfortable export of the computation-heavy ML tree inference to a more powerful computer, faster calculation of diversity statistics, and more as well as easier tree modifications.

As a proof of concept, we obtained the data from two of the the above-mentioned studies: Ding et al. (2020) and Legeay et al. (2020) to reconstruct their phylogenies. To repeat the study by Ding et al (2020), we ran AB12PHYLO with default settings, providing both the raw ab1 files and the sequence data of the type specimens as used by Ding et al (2020). Two samples did not pass the default quality controls. With the remaining 74 samples, we resolved a phylogenetic tree similar to the one in the original work, in which the same genotype groups can be annotated (Figure S2). The MSA used for the phylogeny was 1822 bp long and included 74 samples. Our analysis was run on 12 threads on a system with 64 GB RAM. The total run time for this analysis, including parallelized bootstrapping and BLAST was less than 10 minutes. To repeat the analyses by Legeay et al (2020) we again ran AB12PHYLO with default settings. Again, several samples did not pass the default quality controls. With the remaining samples, we resolved a phylogenetic tree similar to the published one, with a few important differences (Figure S3A), therefore we also remade the phylogeny with the sequence data as deposited on NCBI (Figure S3B) and constructed a phylogeny with all data combined (Figure S3C). These two analyses revealed…

The MSA used for the phylogenies for A, B, C contained 14, 16 and 30 samples respectively, with 2038, 2319 and 2144 sites used for tree inference. On our system, the first two take less than a minute, C around 5 minutes.

Thus, we conclude that AB12PHYLO can produce high-quality multi-gene phylogenies rapidly. The use of AB12PHYLO significantly reduces hands-on working time for these analyses, and overall runtime by parallelization of computation-heavy maximum likelihood tree inference. Moreover, the fact that we observed minor differences between published phylogenies and our re-analyses highlights the importance of reproducible analyses.

## Data Availability

AB12PHYLO is published under the GPLv3 license. It runs on standard desktop computers either under Linux, MacOS or Windows operating systems and can be installed via the pip or conda package-managment systems, the latter also allowing easy installation of an environment with all external tools. Installation instructions and source code are available at https://github.com/lkndl/ab12phylo

## Acknowledgments

We thank Shunping Ding and Marc Buée and colleagues for providing the raw ab1 sequence data from their studies and Tamara Schmey for testing AB12PHYLO.

The project was funded by the German Science Foundation (DFG). Some analyses were performed on the BMBF-funded de.NBI Cloud within the German Network for Bioinformatics Infrastructure (de.NBI) (031A537B, 031A533A, 031A538A, 031A533B, 031A535A, 031A537C, 031A534A, 031A532B).

## Supplementary Materials for

**Figure S1.**
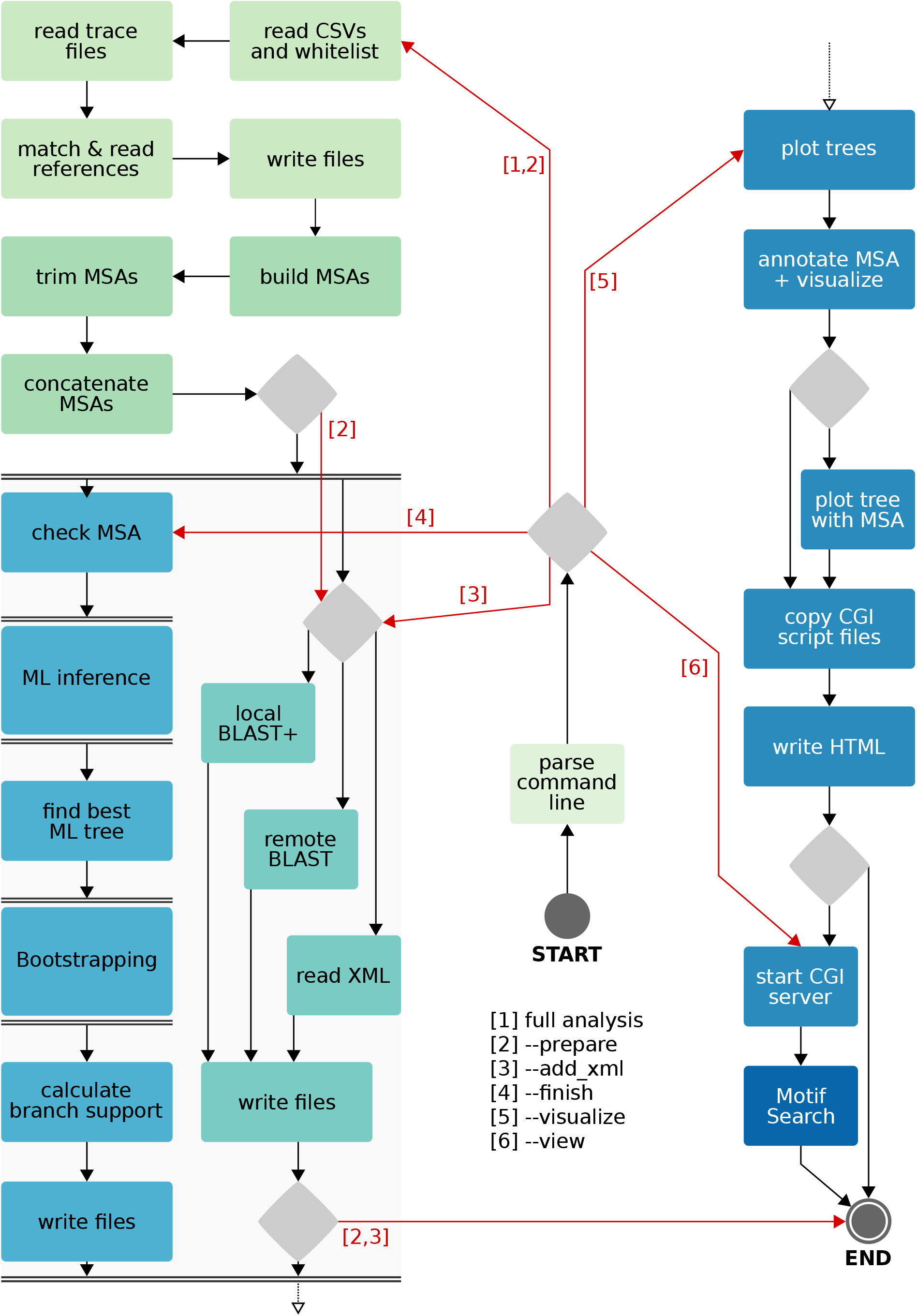
Flowchart of command-line AB12PHYLO. Red arrows represent run modes listed below START, and diamonds mark decisions. The dashed arrow at the bottom signifies that the pipeline continues at the dashed arrow at the top right. Pairs of double horizontal lines indicate the activity between the runs in parallel threads, and the color of the activity rectangles points to the source Python 3 module.

**Figure S2.**
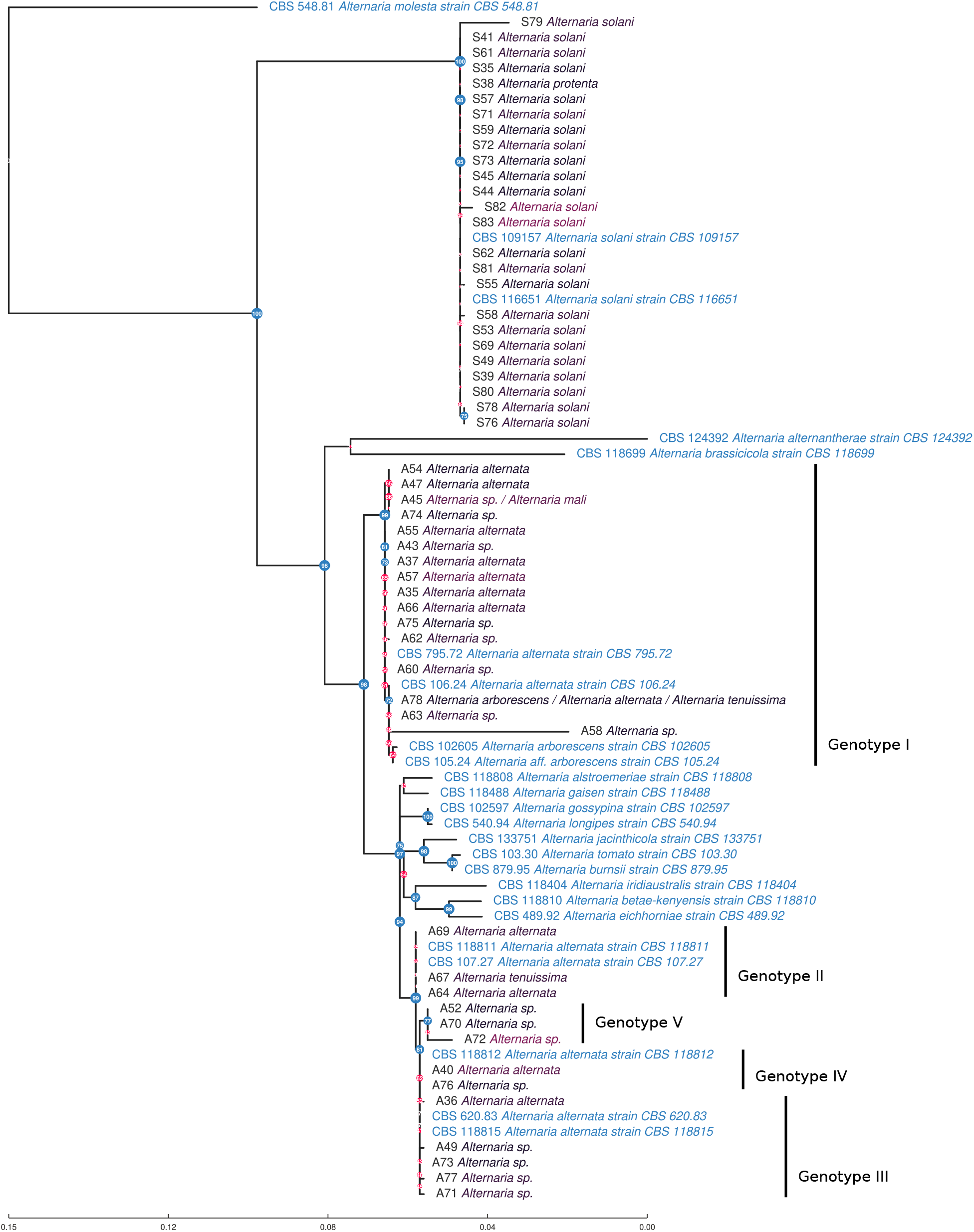
A multi-gene phylogeny constructed with graphical AB12PHYLO from the Alt a 1, EF1, ITS1F and GAPDH loci in the Ding et al. 2020 dataset. The maximum likelihood tree was inferred from 80 ML tree searches, 40 starting at random and 40 at maximum-parsimony trees. The General Time-Reversible (GTR) model of DNA substitution was used in conjunction with four gamma-distributed (+ Γ4) rate classes. Branch support was calculated from 1000 bootstrap iterations and is shown here as Transfer Bootstrap Expectation (TBE, Lemoine et al 2019). Regular samples are labeled with their sample ID and the annotated species of their best BLAST hit, with label color brightening with decreasing percent identity along a black-body spectrum. Reference sequences are labeled in blue. Internal node size and color reflect bootstrap support from 1000 replications, with TBE support > 70% in blue. This is the default style for AB12PHYLO trees.The ML tree was rooted at the reference Alternaria molesta strain CBS 548.81 and agrees with Fig. 3 from Ding et al. 2020. Previously identified genotypes I – V could be resolved and are labeled accordingly.

**Figure S3.**
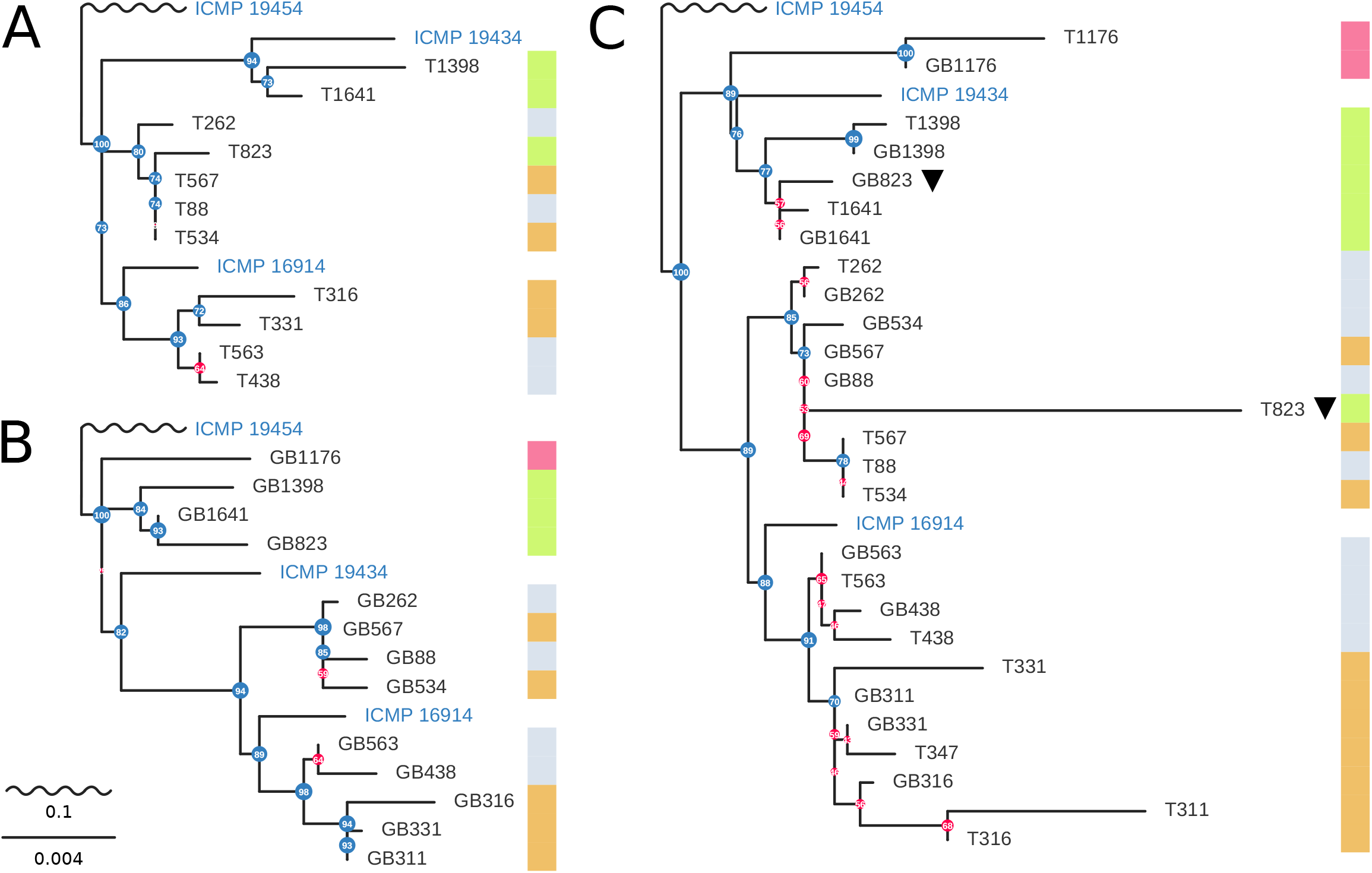
Reconstructions of the phylogeny published in Legeay et al. 2020 based on the ITS, EN1 and YPD-RAS loci, with the colored column on the right indicating which group the sample was originally assigned to. Three NCBI reference isolates for Phytophtora were also included and are labeled in blue. All MSAs were built with MAFFT and trimmed using the balanced Gblocks setting developed for AB12PHYLO. Trees were inferred with RAxML-NG running 240 maximum-likelihood tree searches, from 120 random and 120 parsimony starting trees each. While we used the suggestion from IQ-Tree2 ModelFinder, all three topologies were very robust against changing the evolutionary model to JC, or using IQ-Tree2 instead of RaxML-NG. A: From unpublished ABI trace data, using AB12PHYLO defaults. Note that samples 1176 as well as 311 and 347 were discarded because of low-quality YPD-RAS and EN1 reads. B: Inferred from sequence data submitted to GenBank (entries MH938206–MH938223 and MT598766-MT598818). For sample 347, there is no GenBank record for EN1.C: From both GenBank and ABI trace data. For better reproduction, we built this tree first; adjusting quality control so that trimmed traces visually resembled submitted sequences, and removing more low-quality positions than pre-defined in our balanced Gblocks setting. The colored column on the right indicates the groups previously identified by Legeay et al. 2020.

**Figure S4.**
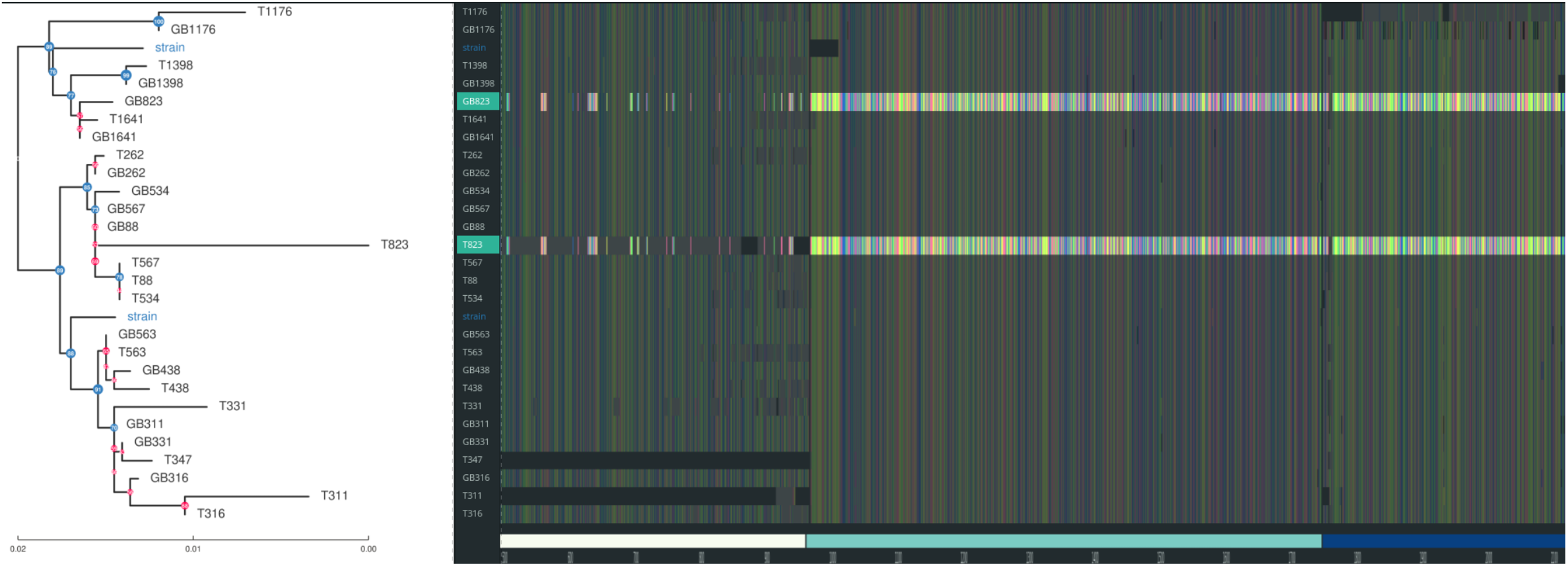
Screenshot of the graphical interface of AB12PHYLO showing the phylogeny from Figure S3C on the left and the quality scores of the samples on the right. The two 823 samples are highlighted. Visible are the large gray and black blocks in T832, the sample derived from the trace file. This indicates that with recommended trimming and QC settings these regions have too low quality to be trusted in the alignment and are therefore omitted. This in turn explain the longer branch length and possibly the discrepancy between the original phylogeny and our reconstruction.

